# A guide to between-community functional dissimilarity measures

**DOI:** 10.1101/2021.01.06.425560

**Authors:** Attila Lengyel, Zoltán Botta-Dukát

## Abstract

One of the effective tools to study the variation between communities is the use of pairwise dissimilarity indices. Besides species as variables, the involvement of trait information provides valuable insight into the functioning of ecosystems. In recent years, a variety of indices have been proposed to quantify functional dissimilarity between communities. These indices follow different approaches to account for between-species similarities in calculating community dissimilarity, yet they all have been proposed as straightforward tools.

In this paper, we review the trait-based dissimilarity indices available in the literature and identify the most important conceptual and technical properties that differentiate among them and that must be considered before their application.

We identify two primary aspects that need to be considered before choosing a functional dissimilarity index. The first one is the way communities are represented in the trait space. The three main types of representations are the typical values, the combination of species×sites and species×trait matrices, and the hypervolumes. The second decision is the concept of dissimilarity to follow, including two options: distances and (lack of) overlaps. We use the above scheme to discuss the available functional dissimilarity indices and evaluate their relations to each other, their capabilities, and accessibility.

## Introduction

Understanding and explaining the variation of living communities along dimensions of space and time have been in the focus of ecological research for a long time. The widely applied scheme by Whittaker (1960, 1972) to tackle questions of different aspects of community variation divides community diversity into alpha (within-community), beta (between-community) and gamma (across-community) components. It is no exaggeration to say that beta diversity sparked the most controversy among these three due to the multitude of ways it can be formulated (Tuomisto 2010a,b, Anderson et al. 2011, Podani & Schmera 2011, Baselga & Leprieur 2015). One of the most popular approaches to beta diversity builds upon the quantification of variation between pairs of communities using dissimilarity indices (Anderson et al. 2006, Legendre & De Cáceres 2013, Ricotta 2017). A broad spectrum of such indices is available for many specific purposes providing elementary tools for different fields of ecology and beyond (see reviews by Legendre & Legendre 1998, Podani 2000).

Nevertheless, choosing from so many options requires a more or less subjective decision from the researcher, which may affect the final result of the analysis. Comparative reviews of dissimilarity indices (Faith et al. 1987, Koleff et al. 2003) and evaluations of the effects of methodological decisions (Lengyel & Podani 2015) help make these decisions.

The most popular, yet not exclusive, interpretations of diversity for a long time considered species as variables that are unrelated to each other. In the last two decades, however, the functional approach to ecological questions gained unprecedented attention (Díaz & Cabido 2001, McGill et al. 2006). This approach relies on the fact that species are not all maximally different from each other; instead, they can be considered more or less related concerning similarities in their traits thought to represent their roles in ecosystems (Violle et al. 2007). The need for explicitly accounting for between-species relatedness generated a wave of methodological improvements that introduced new methods for calculating diversity. Next to a lively scientific discussion on how functional alpha diversity can be appropriately quantified (Mason et al. 2005, Petchey & Gaston 2006, Villéger et al. 2008, Mouchet et al. 2010), some authors developed on the concept of functional beta diversity, too (e.g. Swenson 2011, Botta-Dukát 2018, Chao et al. 2019). Among them, various indices for calculating pairwise functional dissimilarity between communities have been proposed (e.g. Ricotta & Burrascano 2008, Cardoso et al. 2014, Ricotta & Pavoine 2015). Although these indices have been introduced as straightforward measures for revealing between-community dissimilarity based on traits, they have very different concepts behind them, and we still lack a comparative review of them.

This paper aims to provide an overview and a conceptual framework for the pairwise functional dissimilarity measures available in the literature to our best knowledge. We categorise functional dissimilarity indices according to two main properties: (1) the method they represent community composition; (2) the way they define dissimilarity between these representations. We start with a short overview of methods for representing communities based on traits and continue with a discussion of concepts on dissimilarity. Meanwhile, we refer to literature and software sources when appropriate.

### Basic approaches for representing community functional composition

#### Typical value approach

The simplest way to represent the trait composition of a community is to use a single typical trait value for the entire community. In a topological sense, a community is represented by a point in the trait space regardless of the number of dimensions. This approach assumes that within-community trait variation can be neglected (at least, in comparison with the variation between communities). Technically, it takes all individuals within the community as identical. The most widespread example is the community-weighted mean (CWM; Garnier et al. 2004). The rationale behind the CWM can be linked with the mass ratio hypothesis (Grime 1998), stating that the effect of species on ecosystem functioning is proportional to their relative abundances. Although several issues emerged regarding its limited applicability in statistical inference (Hawkins et al. 2017, Peres-Neto et al. 2017, Zeleny 2018) and its negligence of within-community variation (Muscarella & Uriarte 2016, Zheng et al. 2022), CWM is still considered a reliable indicator of trait composition responding to selective forces like environmental matching or succession (De Bello et al. 2007, 2013, Kleyer et al. 2012). For quantitative traits, CWM can be directly estimated from the trait values of individuals within the community or, if individual-level data are unavailable, from the species-level trait values and the (relative) abundances of the species. For qualitative traits, CWM can be defined as the most frequent trait value or as the frequencies of all its possible values. In the former case, CWM is qualitative, similarly to the original trait. In contrast, in the latter case, the available values of the original trait are considered separate variables, and different CWMs can be defined for each. Besides the CWM, other typical values, e.g. the median or the mode, might be considered depending on the scaling of the trait variable and specific research aims. Most frequently, the typical value approach is chosen to simplify a more complicated data structure, for example, when community-weighted means are calculated from species abundances and species-level trait values. However, community-level typical values may be acquired directly for variables of vegetation structure by measuring heights of vegetation layers without considering species composition (Kuchler 1966), for biomass yield by cutting all plant material (’t Mannetje & Jones 2000), or vegetation phenology through remote sensing (Schulp & Alkemade 2011).

### Matrix-based approach

The second approach for representing trait composition is when individuals are not supposed to be identical; however, they can take values only on a discrete scale, typically according to their taxonomical position. This is the usual setting when species abundances are recorded in each community, and each species is assigned a mean trait value (e.g. from external sources). Thus, individuals may (but not necessarily) differ between species but are identical within species. In this case, the species abundances and species-level trait information together represent the functional composition of the community. A common way of calculating between-community dissimilarity building on the above conditions is the use of indices linking a *dissimilarity matrix* between species with a matrix of species abundances. It is especially true if the goal is to calculate functional dissimilarity between not only two but all pairs of a higher number of communities. Let *N* be the number of all communities to be compared and *S* the number of species present in at least one community. The calculation of between-plot pairwise dissimilarities will use an *S*×*S* matrix of between-species dissimilarities and an *N*×*S* matrix of species abundance or occurrence, while the resulting matrix of between-community dissimilarities will have dimensions *N*×*N*. Between-species dissimilarities, *d_ij_*, for species *i* and *j* can be expressed by a measure that satisfies the following conditions: *i* ≠ *j*, 0 ≤ *d_ij_*, *d_ij_* = *d_ji_*, *d_ii_* = 0; moreover, most methods also require *d_ij_* ≤ 1. Dissimilarity indices can be divided into two types: distances and overlaps (de Bello et al. 2013, Mammola 2019). In the context of trait dissimilarity, the most popular distance measure is the Gower index (Gower 1971), for which recently de Bello et al. (2021a) suggested an improved formula. The Gower index takes the (weighted) mean over Manhattan distances between two observations divided by their maximum distances; thus, it ranges between 0 and 1. Since the comparison of functional composition between species and communities does not differ fundamentally in technical terms, any index discussed in the present review that satisfies the above criteria can be adapted for between-species comparisons. Among them, measures of overlap between hypervolumes are especially popular (see *Dissimilarities between hypervolumes* subsection for a more detailed description). However, species-level hypervolumes require the availability of so detailed data about trait distribution that would allow using community-level hypervolumes as well, which are more sophisticated forms of representing community trait composition than the matrix-based approach (see the *Hypervolume approach* subsection).

Another, related form of representing functional composition in a discrete way is the use of a functional classification (Petchey & Gaston 2007), that can be based on external criteria or the original dissimilarity matrix. The classification can consist of mutually exclusive groups of species that can serve as functional types or can express dissimilarities in the form of a dendrogram or a cladogram, in which species are organised in an inclusive hierarchy. This is a widespread approach for accounting for phylogenetic relatedness since phylogenies are commonly summarised in cladograms. Such methods heavily rely on the algorithm chosen for the classification (Podani & Schmera 2006). Examples are provided by Hérault & Honnay (2007), Nipperess et al. (2010), and Cardoso et al. (2014). However, while phylogenies inherently assume tree-like relationships of species through common ancestors, such models have no conceptual support in the case of traits. Due to this reason, and since recently, excellent reviews have been published on phylogenetic beta diversity (Pavoine 2016, Tucker et al. 2017), we omit classification-based indices in the present paper. Nevertheless, all indices we review can be freely used also for quantifying phylogenetic dissimilarity if between-species relatedness represents phylogenetic distance.

### Hypervolume approach

The third form of representing communities in the trait space accounts for between-individual variation on a continuous scale. This is done either by defining a single continuous probabilistic distribution for the community, or by combining separate distributions defined for each species, or even for lower units (e.g. age classes) into one community-level distribution (Carmona et al. 2016, 2019). Describing communities by distributions means assigning probabilities to the points of the trait space, referring to the likelihood of being occupied by an individual from the community (Blonder et al. 2014). In more practical terms, it means delineating occupied and unoccupied parts of the trait space with sharp or fuzzy boundaries. A traditional way to differentiate between occupied and unoccupied points is the use of hypervolumes. Hypervolumes were introduced initially by Hutchinson (1957) to map the fundamental niche of species in a multidimensional environmental space. By considering trait dimensions as niche axes, we can adopt this methodology to represent trait distributions of communities. However, delineating hypervolumes is challenging, and several conceptually different solutions are available. Blonder (2018) has recently provided an in-depth review of hypervolume concepts; hence we discuss them here only briefly. Notably, some types of hypervolumes are designed explicitly for comparing populations or species and are not adapted for communities (e.g. MacArthur & Levins 1967, Mouillot et al. 2005, Geange et al. 2011, Jarvis et al. 2019, Lu et al. 2021).

Hutchinson (1957) used multivariate range boxes (MRB) as hypervolumes. Range boxes are finite sections of the multidimensional space defined by minimum and maximum values of ’occupied’ points along each axis independently. Convex hulls are convex sets that enclose all occupied points within the smallest volume (Cornwell et al. 2006). MRBs and convex hulls do not differentiate between points included; thus, they are insensitive to within-set variation in the density of points; practically, they utilise only presence/absence information. Thanks to this, they are compatible with matrix-type data, too (see the *Matrix-based approach*).

Hereafter we call them *unweighted hypervolumes*, in contrast to *weighted hypervolumes* that differentiate between the points within the boundaries reflecting the likelihood of being occupied. The first example of weighted hypervolumes is the dynamic range boxes (DRB) suggested by Junker et al. (2016). Range boxes can be shrunk in a way to contain only the narrowest α (0 < α < 1) proportion of the data according to the quantiles of the empirical distribution, starting with the entire interval and ending with only the median. With changing α, a sequence of MRBs can be obtained. We can calculate a volume for each MRB, and averaging them provides a parameter-free estimation of the hypervolume size. To partial out eventual between-trait correlations, it is advisable to transform trait variables into principal components. The probabilistic nature of hypervolumes can be taken into account also if they are defined by modelling techniques with continuous outcomes. Such methods include fitting multivariate normal distributions, kernel density functions, linear, non-linear or tree-based regression methods, or machine learning classifiers. They all provide a hypervolume function that can be interpreted as a probability density function, although with different assumptions and constraints. It is notable that despite these functions do not provide, by definition, sharp boundaries for the hypervolume, for simplifying calculation, they are often discretised at a low cut-off threshold (Carmona et al. 2019).

Notably, hypervolumes can be fitted on points in the trait space (e.g. typical values at the level of communities or species) even if individual-level trait values are not available. In this case, one needs additional assumptions or information about the parameters of the hypervolumes (Carmona et al. 2019).

### Concepts and methods for calculating functional dissimilarity between communities

#### Distances and overlaps

The concept of functional dissimilarity must be formalised in different ways according to how we have defined functional composition at the community level (Fig. 1). The indices currently available in the literature mirror two main, non-exclusive concepts of dissimilarity: *distances* and *(lack of) overlaps* (for simplicity, hereafter, we refer to the latter as overlaps). Distances express how far two communities are from each other in the trait space; alternatively, the degree to one community’s position needs to shift until it becomes identical to the other community. In the presence of several variables, these indices summarise differences in the values of each variable to calculate a multivariate distance. A common example, Euclidean distance takes the square-root of the sum of squared differences, while Manhattan distance is the sum of the differences in each variable between the two communities. These methods are sensitive to the scaling of the variables because those with higher variation contribute more to the final distance. Thus in many cases, it is straightforward to standardise them; however, after standardisation, they may obtain an interpretation also as overlap-type indices.

**Figure 1.**
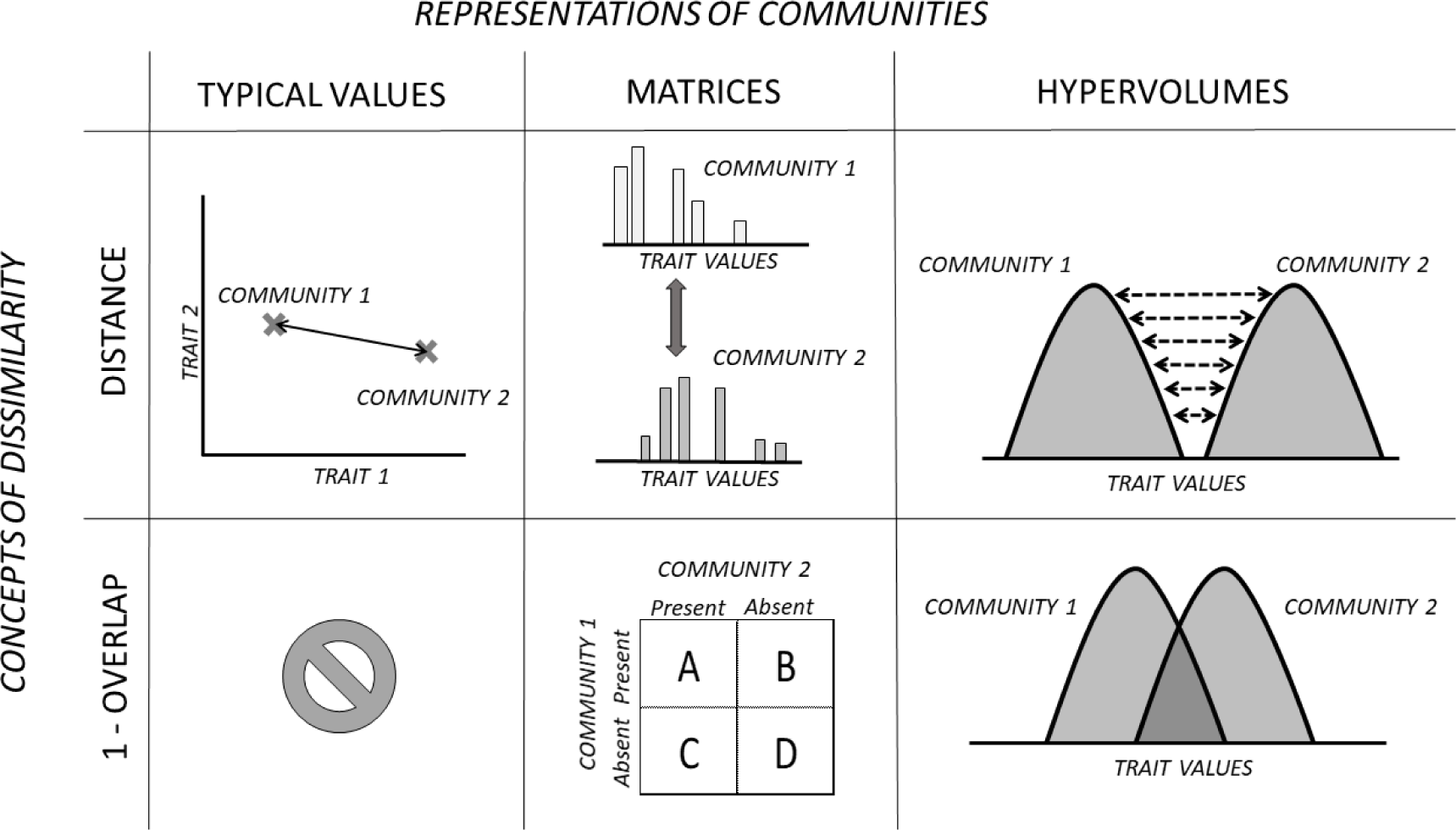
A scheme of the classification of functional dissimilarity measures. The two fundamental decisions are the representation of communities in the trait space and the concept of dissimilarity to be applied. There is no available method for measuring overlap of typical values, since points have no extent.

Overlaps decompose the diversity of two communities into shared and unshared quantities to build up an index from them. Most overlap indices range between 0 and 1 and can be written in either similarity (*s*) or dissimilarity (*d* = 1 – *s*) form. (Hereafter, when we see it unnecessary to differentiate between similarities and dissimilarities, we use the term *resemblances*.) Such indices are popular in species-based analyses; thus, we follow these primers to introduce them. In the case of presence/absence data, any resemblance can be calculated from four quantities (denoted by *a*, *b*, *c*, and *d*, respectively): the number of shared species, number of species occurring only in the first or the second community, and species missing from both communities; however, the last one is often neglected (but see Tamás et al. 2001). The most widely used indices calculate similarity by dividing the number of shared species by its upper limit. Such limit may be the arithmetic, geometric or harmonic mean or the minimum of species richness in the two compared plots. All of these indices range from zero to one. Note that quantity *a* can be interpreted as shared diversity, while *a+b* and *a+c* as the diversities of the two compared communities. With presence/absence data of species composition, the most popular indices include the Jaccard and Sørensen indices (*S_Jaccard_* = *a/(a+b+c)*, *S_Sørensen_* = *2a/(2a+b+c)* in similarity form; Jaccard 1901, Sørensen 1948), and many of the alternatives are algebraically related to them. For abundance data, the dissimilarity of two communities can be derived from the summation of species-wise differences, thus revealing a direct link between overlaps and distance-type indices. The theoretical maximum of Euclidean and Manhattan distances (the highest possible distance between two communities) is the species’ total abundances, making them difficult to compare across samples; therefore, indices involving standardisation have become more favoured in ecological studies. Standardisation is possible through standardising species-level differences and then summing or averaging them, as done in the Canberra metric (Lance &Williams 1966) and two versions of the normalised Canberra metric (Ricotta & Podani 2017). Another way of standardisation is first summing raw species-level differences and then dividing this sum by the sum of their upper limits. The upper limits can be approached by the sum or maximum of species’ abundances, resulting in Bray-Curtis (Bray & Curtis 1957) and Marczewski-Steinhaus (Marczewski & Steinhaus 1958) indices, respectively. For presence/absence data, they reduce to Sørensen and Jaccard indices, respectively. More extensive reviews of resemblance indices are available in Hubálek (1982), Legendre & Legendre (1998), Podani (2000), Koleff et al. (2003), while Ricotta & Podani (2017) explains several implicit relations between indices. See also Appendix A1 for the summary of indices mentioned in this paper. Functional dissimilarities designed analogously with the species-based overlap indices introduce alternative ways to calculate shared diversity and diversities of compared communities and replace *a*, *a+b* and *b+c* quantities with new, trait-based measures.

Species-based overlaps in dissimilarity form reach their maximum (that is, unity) when the two compared communities do not share any species. In this context, we could call such communities *maximally distinct*. However, when traits are considered, two communities can be similar, even if they do not share any species. For example, suppose a similar species in community B replace all species in community A. In that case, the two communities have no shared species, but they are similar from the functional point of view. In the functional context, two communities are maximally distinct when the similarity of any species from the first community is zero to any species in the other community. It is a desirable property for a functional dissimilarity index to take a maximum value if and only if the two compared communities are maximally distinct (Botta-Dukát 2018). Notably, this also implies the involvement of between-species dissimilarity in a similarly bounded form that we can obtain by Gower distance or an overlap-type dissimilarity.

The framework described below is summarised on Table 1.

**Table 1.**
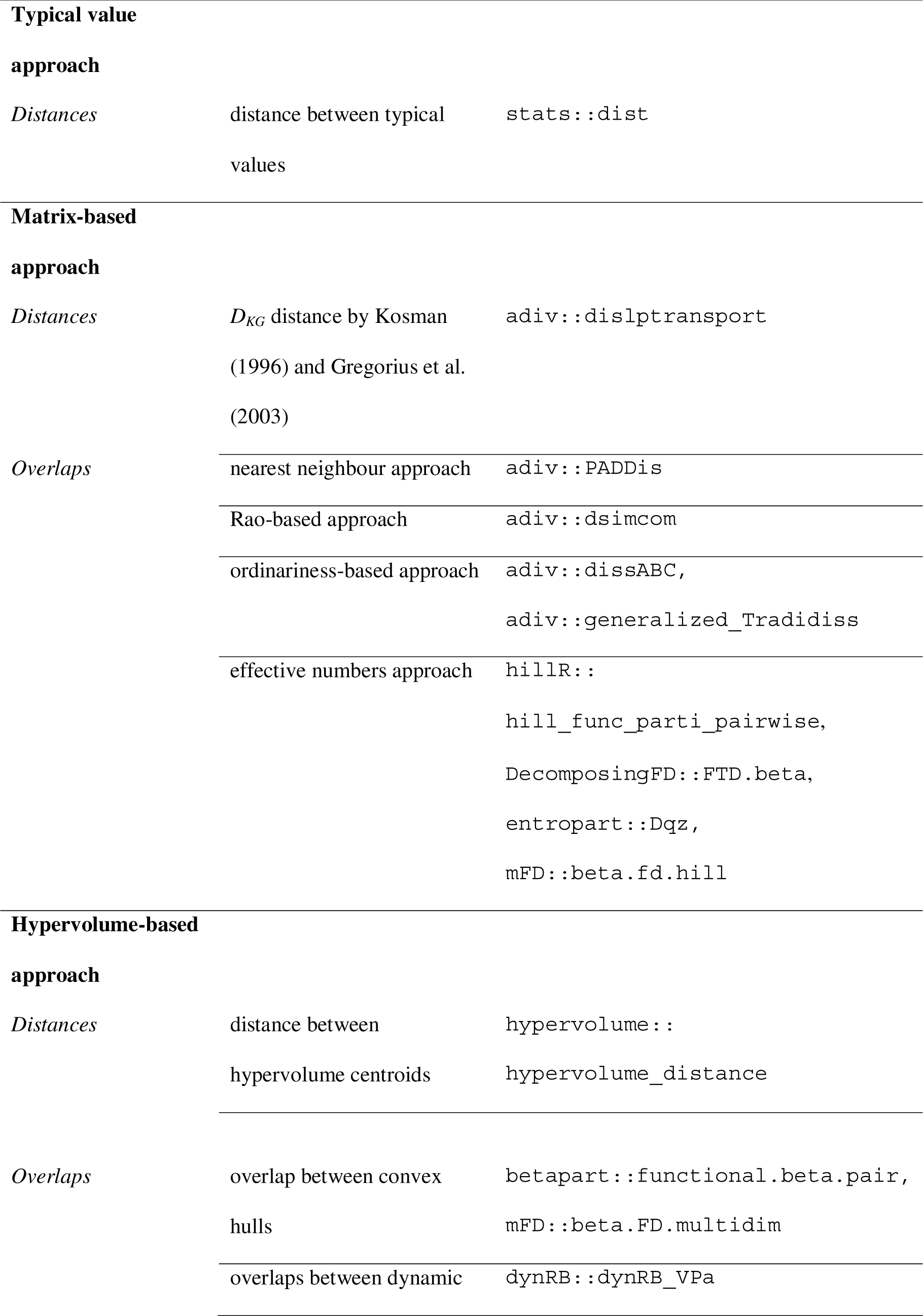
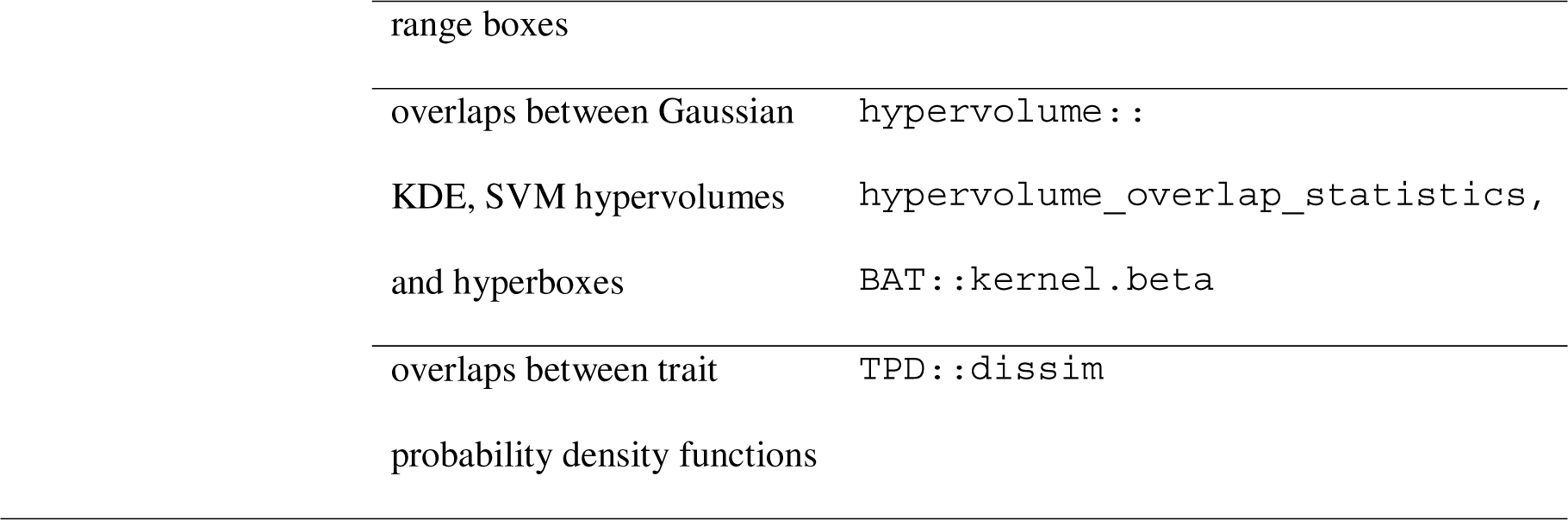
The scheme of classification of between-community functional dissimilarity indices with available R functions

#### Dissimilarities between typical values

If communities are represented as typical values in the trait space, then distances are the only option for calculating dissimilarity. The Euclidean or Manhattan distance between CWMs (hereafter called *CWMdis*) is a straightforward way of expressing between-community dissimilarity. One of its merits is the simple calculation that makes it easy to analyse data matrices comprising thousands of communities and species within a feasible runtime and capacity demand (Lengyel et al. 2020). Another advantage of this method is its Euclidean property; thus, their interpretation is close to our perception of physical distance. For qualitative traits, *CWMdis* can take the values 0 (agreement) or 1 (mismatch). Since CWM is commonly used for representing the trait composition of communities, the Euclidean distance between CWMs is a natural way for comparing communities, either explicitly (e.g. Müller et al. 2021, Engbersen et al. 2022) or implicitly as part of a linear model (e.g. Moretti et al. 2013, Prieto et al. 2017). Ricotta et al. (2015) investigated the relatedness of the distance between CWMs (hereafter called *CWMdis*) with the Rao-based approach (see therein) and showed its applicability also on phylogenetic data. Sometimes more complicated representations are also simplified to a typical value for calculating dissimilarity between them. For example, a possible way of comparing hypervolumes is measuring the distance between their centroids, their closest points, or specified quantiles (Blonder et al. 2018, Mammola 2019); however, with typical values other than the centroids, the Euclidean property of a between-community distance matrix is unlikely to hold.

#### Dissimilarities between matrix-type representations

Among distance-type indices, there are few options to choose from if community trait composition is represented in a matrix form. Kosman (1996) and Gregorius et al. (2003) proposed an algorithmic index, hereafter called *D_KG_*, for the quantification of differences between distributions of numerical traits interpreted as discrete variables. *D_KG_* is the minimal sum of frequencies shifted from one trait value to another, weighted by the magnitude of differences between the respective values. Minimising the sum of shifted frequencies when approaching from one distribution to another is known in linear programming as the transportation problem (Hitchcock 1941). This index was recently re-iterated and implemented by Ricotta et al. (2021); apart from its introductory publications, it has not yet been applied in case studies. If species are grouped into functional types, then it is also possible to analyse the [communities] × [functional groups] matrix by any dissimilarity measure compatible with species data (e.g. Euclidean or Manhattan distance, but overlap indices as well). Hérault & Honnay (2007) presents an example of such application.

Scheiner et al. (2017) proposed measuring the functional difference between two communities, *A* and *B*, by the mean of mean distances of species occurring in the first community from species occurring in the second one (*m_AB_*). However, this index can take a positive value even if two identical communities are compared, thus an adjustment is for obtaining a straightforward distance measure:

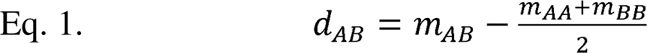

Kosman (2014) and Kosman and Leonard (2007) proposed a similar measure, but based on average distances between individuals instead of species, calling it “distance of average differences”. If intraspecific trait variation is not considered, mean distances between individuals can be calculated as the mean distance between species weighted by the products of their relative abundances, including the zero distance of species from itself. In this case, *d_AB_* is equivalent to Rao’s dissimilarity (Rao 1982), also called *DISC* (Pavoine & Ricotta 2014). Kosman (2014) and Kosman and Leonard (2007) cautioned that it is not guaranteed that this measure is always non-negative. Fortunately, Rao’s dissimilarity is non-negative for dissimilarities with Euclidean property (Pavoine et al. 2005).

Overlap indices offer a diverse toolbox for calculating functional dissimilarity between matrix-type representations. Here we classify and introduce them according to the concepts they follow.

### Nearest neighbour indices

According to this approach, the resemblance of two communities is determined by the most similar pairs of species between the two communities. Looking at species as maximally different and taking *X* and *Y* the two communities under comparison, *b* quantity of the 2×2 contingency table can be viewed as the total uniqueness of community *X*. The uniqueness of a single species in *X* is 1 if it is absent in *Y*; otherwise, it is 0. Therefore, *b* is the sum of the species’ uniqueness values. However, from a functional perspective, the uniqueness of a species present only in *X* should be between 0 and 1 if it is absent in *Y,* but a similar species is present there. Therefore, it is possible to define the analogue of *b,* which accounts for similarities between species:

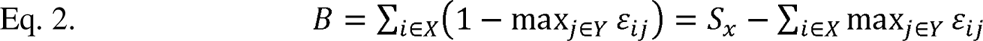

where species in *X* are denoted by *i*, species in *Y* denoted by *j*, ε*_ij_* is the similarity between species *i* and *j*, *S_X_* and *S_Y_* are the species richness for *X* and *Y*, respectively. The same logic applies to *c*, which is the uniqueness of community *Y*, where *C* expresses the degree of functional uniqueness:

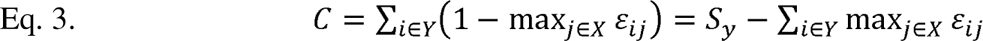

Ricotta et al. (2016) define shared diversity (*A)* as follows:

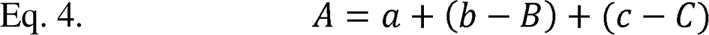

Having *A*, *B*, and *C* defined as analogues of *a*, *b*, and *c*, it is now possible to design trait-based similarity measures following the logics of Jaccard, Sørensen, Sokal & Sneath (1963), Kulczynski (1927), Ochiai (1957) and Simpson (1960) indices resulting in a new, general family called *PADDis* (Ricotta et al. 2016). Notably, Ricotta et al. (2016) define *A* as a quantity that ensures the components A, *B* and *C* to add up to *a + b + c* but with no explicit biological interpretation.

The earliest representatives of this approach were shown by Clarke & Warwick (1998) and Izsák & Prince (2001). Ricotta & Burrascano (2008) and Ricotta & Bacaro (2010) developed the *D_CW_* and *D_IP_* indices, which are identical with the Sørensen and Kulczynski forms of *PADDis*. Although *PADDis* was primarily defined for the presence-absence data type, the generalisation of *D_IP_* and *D_CW_* to relative abundances, *D_CW_(Q)*, was also derived by Ricotta & Bacaro (2010). For these two versions, it is not necessary to explicitly define the *A* component. Using the relationships between Jaccard, Sørensen, Kulczynski, Ochiai and Sokal & Sneath indices (Appendix A2), it is theoretically possible to derive the extension of *PADDis* to relative abundances from *D_CW_(Q)*; however, the biological interpretation of *A* remains dubious in this framework. In general, the nearest neighbour-based indices are moderately popular compared with the other dissimilarity indices mentioned above. However, they were applied more frequently in phylogenetical than in functional contexts, e.g. Greenlaw et al. (2011), Ricotta et al. (2012), but see Sonnier et al. (2014).

The differential feature of the nearest neighbour approach is that it is based on the dissimilarities between species in one community and their most similar species in the other community. In contrast, the following approaches are based on an average dissimilarity between species in one community and all species in the other.

### Rao-based indices

This family of indices can be traced back to the diversity framework proposed by Rao (1982) and recently extended by Pavoine & Ricotta (2014). Now a wealthy group of methods builds upon this concept and provides several examples of crossing the boundary between distances and overlaps; however, as its most promising members are overlaps, we discuss them here. Rao’s within-community diversity is defined as the expected dissimilarity between two randomly drawn individuals from a single community [*Q*(**p**)]. This has become a widely used index of functional alpha diversity (Botta-Dukát 2005). Likewise, a between-community component of diversity [*Q*(**p**,**q**)] can be defined as the dissimilarity between two random individuals, each selected from different communities. Subtracting the mean within community diversity from the between community diversity leads to Rao’s dissimilarity (also called *DISC*). If dissimilarity between species ranges from zero to one and has Euclidean property, 0 ≤ *DISC* ≤ 1 (Pavoine & Ricotta 2014). However, *DISC* may be much less than 1, even if the two communities are completely distinct. Therefore, Pavoine & Ricotta (2014) suggested dividing *DISC* by its theoretical maximum (see equations 3 and 4 in Pavoine & Ricotta 2014). They recognised that the resulting index is a representative of a broader family of indices, hereafter called *dsimcom*, which are actually the implementations of Rao’s between-community and within-community components of diversity into the similarity formulae designed for presence/absence data. For this family of indices, the expected similarity between individuals of different communities is taken as analogous with the shared diversity (*a*), while the expected similarities within communities are analogous with the diversity (species numbers) of two communities (*a+b*, *a+c*). In this way, Pavoine & Ricotta (2014) presented formulae following the Sokal & Sneath (1963), Jaccard, Sørensen, and Ochiai indices. Pavoine & Ricotta (2014) showed that members of the *dsimcom* family provide meaningful values also if absolute abundances or presence/absence data are used instead of relative abundances. While the sum of relative abundances in a community is always one, two communities may differ in the sum of absolute abundances (total abundance) and the sum of presences (species richness). When absolute abundances or occurrences replace relative abundances, these differences also contribute to dissimilarity. An advantage of this family is that if the between-species similarity matrix is positive semi-definite, the matrix of between-community similarities is also positive semi-definite (Pavoine & Ricotta 2014). It implies that the distance matrix calculated as the square-root of the complements of similarities has Euclidean property (i.e. embedded into a Euclidean space). This makes the interpretation and visualisation of multivariate patterns revealed by these indices more straightforward. When the between-species similarity matrix contains taxonomical similarities, its off-diagonal elements are 0, and for presence/absence data *A*=*a*, *B*=*b*, and *C*=*c*.

It is worth noting the inherent link between *DISC* and *CWMdis* based on the geometric interpretation by Pavoine (2012) and Ricotta et al. (2015). Pavoine (2012) showed that if between-species dissimilarities are in the form *δ_ij_* = (*d_ij_*^2^)/2 and *d_ij_* has Euclidean property, *DISC* is half the squared Euclidean distance between the centroids of two communities – a function monotonically related with *CWMdis*, the simple Euclidean distance between centroids of communities. As Ricotta et al. (2015) argue, if species relatedness is only described by a dissimilarity matrix, which is the common case in phylogenetic analyses, species can be mapped into a principal coordinate analysis ordination using *d_ij_*. Given the Euclidean embeddable property of *d_ij_*, this ordination should produce *S*-1 or fewer ordination axes, all with positive eigenvalues. Ordination scores for species can be used as traits; therefore, centroids of communities and (squared) Euclidean distances between communities can be calculated. In the particular case when between-species dissimilarities are Euclidean distances, *DISC* must be equal to the Euclidean distance between the weighted averages of traits, that is, *CWMdis*.

It is also notable that Swenson et al. (2011) and Swenson (2011) use the quantity *Q*(**p**, **q**) as a standalone index of pairwise beta diversity and call it *D_pw_* or “Rao’s *D*”. The latter name is misleading since Rao (1982) himself noted with *D_ij_* the *DISC* (or *D_Q_*) index. *Q*(**p**, **q**) measures dissimilarity between two communities but the dissimilarity of a community from itself is not zero. Swenson (2011) also presents a standardised version of *Q*(**p**, **q**) under the name “Rao’s” H. With this formula, the dissimilarity of a community to itself is scaled to 1, and its transformation to a meaningful scale where each community has dissimilarity value zero towards itself is not elaborated.

Schmidt et al. (2017) proposed probabilistic indices with weighted and unweighted versions for expressing community similarity based on taxa interaction networks (called *TINA*, taxa interaction-adjusted) and phylogenetic relatedness (*PINA*, phylogenetic interaction-adjusted). *TINA* and *PINA* differ only in what type of data the interaction matrix contains. Notably, the functional formula of weighted *TINA* is identical to the Ochiai version of *dsimcom*. However, the unweighted *TINA*, abbreviated *TU*, is not a special case of *TINA*, which we consider an inconsistency.

Until now indices following the Rao-based approach have been applied sporadically (e.g. Díaz-Varela et al. 2016, Sobrinho et al. 2016, Sfair et al. 2016).

### Ordinariness-based indices

Concerning functional alpha diversity, Leinster & Cobbold (2012) introduced the concept of species ordinariness, defined as the weighted sum of relative abundances of species similar to a focal species within the same community, or in other words, the expected similarity of an individual of the focal species and an individual chosen randomly from the same community.

According to Ricotta & Pavoine (2015), it is straightforward to replace abundances with ordinariness values in the species-based (dis-)similarity indices. Following this concept, they introduced a new family of trait-based similarity measures called *dissABC*. *dissABC* is based on Tamás et al.’s (2001) generalisation of presence/absence indices but replaces abundances by ordinariness. Ordinariness values can be calculated as the weighted sum of relative or absolute abundances, either with respect to the pooled species list of the two communities under comparison or to the total species list of the data matrix containing other communities.

According to Pavoine & Ricotta (2019), by replacing species abundances with species ordinariness values in the generalised (i.e. weighted) Canberra index, a meaningful dissimilarity index can be designed, which is called *generalized_Tradidiss*. The generalised Canberra index weighs the contribution of each species to the overall dissimilarity between the two communities. We can set this weight to be constant for all species (leading to an analogue of the normalised Canberra index) or weigh them proportionally to their relative abundance in the pooled communities (obtaining an analogue of the Bray-Curtis index). The generalised Canberra index standardises species-level differences by the sum of species abundances in the two communities. However, standardisation is also possible by the maximum instead of the sum, thus leading to analogues of the Marczewski-Steinhaus index.

Ricotta (2018) has proposed a dissimilarity index based on the evenness of species abundances. This index can be calculated not only for abundances but also for ordinariness, thus leading to another functional dissimilarity index accessible in the *generalized_Tradidiss* framework.

More recently, Ricotta et al. (2022) proposed another family of indices, where the importance of large differences in ordinariness compared to small ones can be adjusted following the Minkowski distance framework.

The functions *dissABC* and *generalized_Tradidiss* are available in the adiv R package (Pavoine 2020). We are unaware of any case studies applying the ordinariness-based indices mentioned above.

### Effective numbers indices

When interspecific similarities are not considered, there is emerging consensus that using effective numbers (also called number of equivalents or Hill diversity) is a straightforward way for partitioning diversity into within-community (alpha, *<ι>α*), and between-community (beta, *<ι>β*) components (MacArthur 1965, Hill 1973, Jost 2007, de Bello et al. 2010). Effective numbers have a parameter called the *order of diversity* (*q*) that allows setting the weight given to rare species, e.g. *q* = 0 representing the presence/absence case, *q* = ∞ considering only the relative abundance of the most abundant species in the community. The main advantage of using an effective number is that the magnitude of alpha and gamma diversity is expressed in units of species, similarly to the number of species, which is its *q* = 0 case. Besides easier interpretation, this provides independent alpha and beta components from multiplicative diversity partitioning. Multiplicative beta diversity (*<ι>β*_multi_=*<ι>γ/a*) in effective numbers is the number of maximally distinct communities with the observed alpha diversity that would results in the observed gamma diversity. Therefore, beta diversity ranges from 1 (all communities are the same) to the number of communities (all of them are maximally distinct), that is, to 2 in the case of pairwise comparisons. For a more convenient use, *<ι>β*_multi_ should be rescaled to the interval [0; 1]. Earlier works (Harrison et al. 1992, Jost 2006, Chao et al. 2012) suggested two approaches to convert the ratio of alpha and gamma diversities into pairwise dissimilarity:

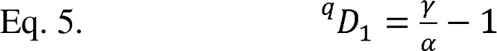

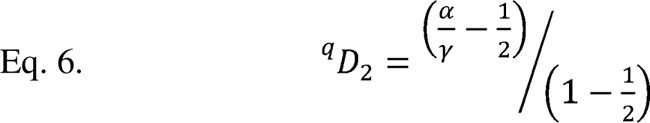

In the lack of agreement on the naming of the above indices, we follow Chao et al. (2012) in calling *^q^D_1_* “turnover”, while we use the term “true overlap” (see Wolda 1981, Jost 2006) for *^q^D_2_*. Turnover is a linear function of *<ι>β*_multi_, while true overlap relates linearly to the inverse of *<ι>β*_multi_. With presence/absence data (*q* = 0), overlap reduces to the Sørensen index, with *q* approaching 1, it is equivalent to Horn index (Horn 1966), and for *q* = 2 it reduces to the Morisita-Horn index (Horn 1966, Jost 2006). The two rescaling formulae provide strongly correlated results.

The implementation of functional similarity of species into the framework of the effective numbers inspired several attempts (Leinster & Cobbold 2011, Chiu & Chao 2014, Scheiner 2012, Scheiner et al. 2017). Leinster and Cobbold (2011) introduced the concept of species ordinariness, the expected similarity of an individual of a species in one community to a randomly chosen individual in the other community. The ordinariness of a community can be calculated by taking the average of species ordinariness values weighted by relative abundances, while the reciprocal of the average ordinariness gives the effective numbers of species. By changing the *q* value, it is possible to weigh the influence of dominant species in the calculation of the average using the formula of the generalised mean (Hardy et al. 1952). This framework is the most popular currently (e.g. Veresoglou et al. 2014, Abrams et al. 2021). Effective numbers of species according to Leinster & Cobbold can be easily partitioned into alpha, and beta components in the multiplicative way. Then, this multiplicative beta can be rescaled using the formulae mentioned above. These indices behave consistently only if abundances are taken into account as relative abundances.

Besides Leinster & Cobbold (2011), several authors suggested alternative formulations of effective numbers based on functional dissimilarity of species within a community. Scheiner (2012) derived such a metric from the evenness of nearest neighbour distances between species, while the approach of Presley et al. (2014) relies on the total interspecific distances. Both these approaches stress the importance of separating the effects of varying abundance and functional dispersion of species. However, these metrics are not elaborated for deriving between-community dissimilarity. Building on the evenness of pairwise dissimilarities of communities, Scheiner et al. (2017) defined *<ι>β* diversity using the evenness of pairwise distances between communities combined with mean dispersion. The resulting index (denoted as *^q^D(TM)*) is not derived directly from γ and α diversity; however, it also takes the value *N* if the *N* communities are completely distinct, and equals 1 if they are identical. In the approach proposed by Chiu and Chao (2014) two communities are maximally distinct when they have no shared species. Therefore, their beta diversity may be maximal even if a very similar species replace each species.

The concepts of multiplicative and additive diversity partitioning with Hill’s numbers are increasingly known in functional diversity; however, we are unaware of any application as a pairwise dissimilarity measure. The methods mentioned above are available from the R packages entropart (Marcon & Herault 2015), hillR (Li 2018), mFD (Magneville et al. 2022), and Decomposing FD (by S. Kothari, https://github.com/ShanKothari/DecomposingFD).

### Dissimilarities between hypervolumes

When expressing dissimilarity between hypervolumes, we face the same dichotomy as earlier: we can choose the distance or the (lack of) overlap between hypervolumes. According to Mammola (2019), these two are complementary aspects of between-community dissimilarity, and the best practice is applying both. From a theoretical point of view both distances and overlaps can be defined for each variant of hypervolumes; however, hypervolume representation methods vary in which forms of dissimilarities are available for them in current software.

Among distances, a simple and popular index is the distance between centroids of hypervolumes (Blonder et al. 2014, 2018), which leads back to approaches when communities are represented by typical values (Fig. 1). Actually, the difference between the CWM and centroid lays in that CWM is typically calculated from the raw data, while the centroid is usually estimated from the fitted hypervolume. In case of a hypervolume fitted perfectly on the data, there should be no difference between them. Obviously, in this case the information on within-community variation, the main advantage of using hypervolumes, is lost. See the section “Typical values” for details. Similarly to *CWMdis*, the distance between centroids is a robust measure of communities differentiated along trait gradients (Fig. 2a-c). However, this method does not indicate difference between communities that differ only in within-community trait variation (Fig. 2d-e). It is also possible to measure distance between the closest points of a pair of distributions. However, minima and maxima are often not robust, thus the truncation of the hypervolumes at certain quantiles might be straightforward. For example, by applying a 5% threshold, in each hypervolume the most distant 5% of points in all directions are disregarded. Notably, there are two significant drawbacks of applying quantiles instead of the centroid to calculate distance between hypervolumes. The first one is that when calculating distance between all pairs of a set of communities, in each pairwise comparison different points of the hypervolumes are used as reference for the same hypervolume. This may result in a non-Euclidean distance matrix limiting possibilities of subsequent analyses. The second drawback is that if two hypervolumes overlap (even after truncation), then distance between quantiles cannot be defined on non-negative range, or it will not be sensitive to different levels of overlap.

**Figure 2.**
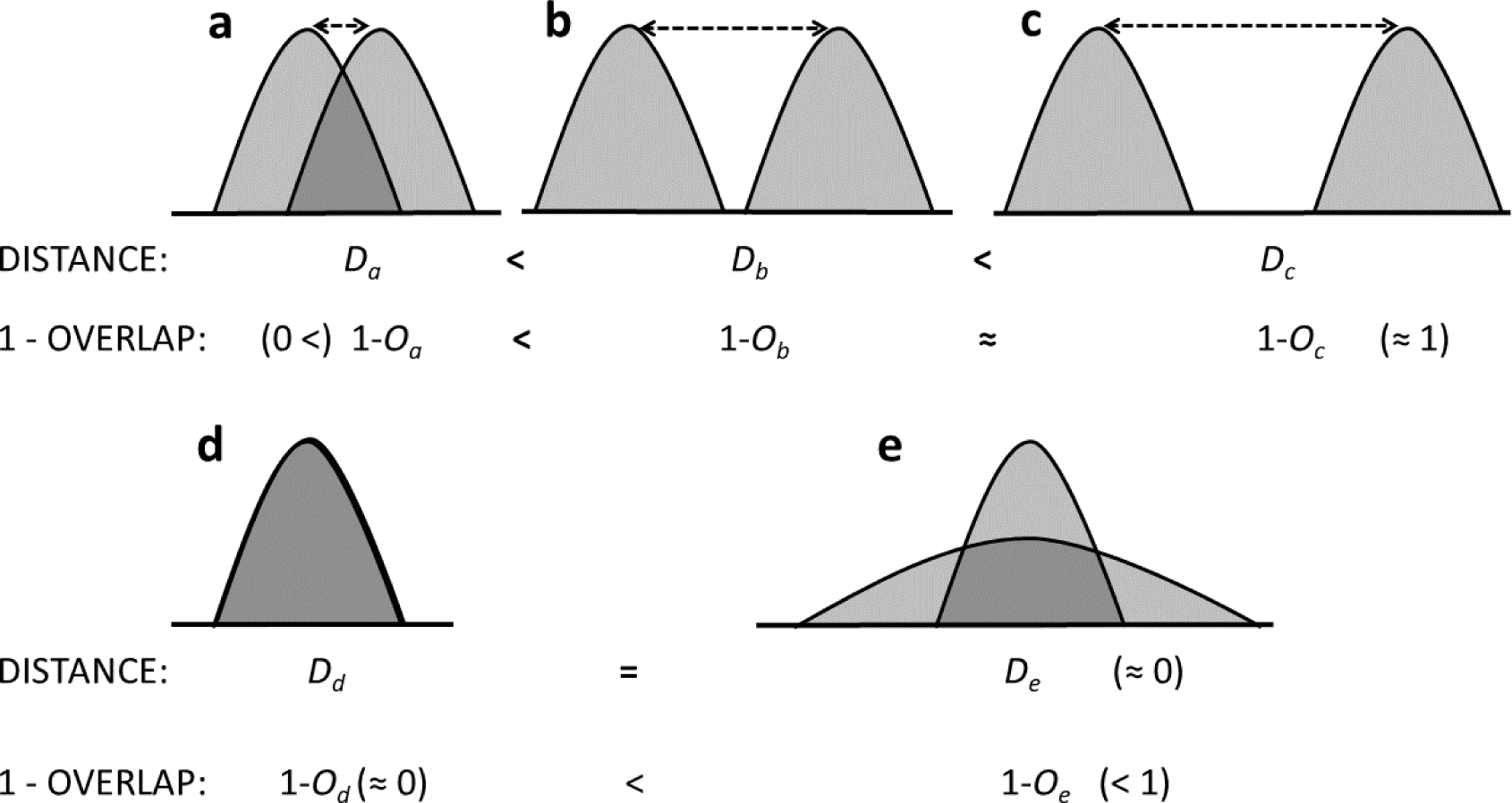
Behaviour of distance-type and overlap-type (1-overlap) dissimilarities under different relations of hypervolumes. a) the two hypervolumes have some overlap, and there is some distance between their centroids; b) hypervolumes have no overlap, their centroids are further apart than at a); c) no overlap between hypervolumes, centroids are even further apart, than at b); d) the two hypervolumes maximally overlap, centroids are identical; e) the centroids of the two hypervolumes are identical but their overlap is not maximal due to different variance of hypervolumes.

Overlap can be interpreted as the shared portion of diversity and can be calculated either in absolute terms or can be standardised by dividing it by the summed or the mean size of the hypervolumes, leading to analogues of the Jaccard and the Sorensen index (e.g. Villéger et al. 2011, 2013), respectively. Such indices are already available for convex hulls, hypervolumes based on Gaussian kernels and support vector machines (Baselga et al. 2022, Blonder et al. 2022), and trait probability density functions (Carmona 2019). An alternative method to compare hypervolumes *h_1_* and *h_2_* is to divide the absolute overlap by the total volume of *h_1_*, leading to an asymmetric index between *h_1_* and *h_2_*, and vice versa. The mean of the two asymmetric indices can be used as a symmetric measure, as it is already available for dynamic range boxes (Schreyer et al. 2022). One of the merits of overlap indices is their sensitivity to the shape and size of the hypervolumes. Even if the centroids of the two communities in the trait space are identical, there might be difference between them in the extent the points are spread around the centroid (Fig. 2d-e). Such differences are usually detectable using overlap measures. On the other hand, the disadvantage of overlap indices is that if two hypervolumes have no overlap, it is indifferent how far they are from each other in the trait space, their dissimilarity is (approximately) constantly maximal (Fig. 2b-c). That is, these indices are especially straightforward in the comparison of communities not completely separated in the trait space (Fig. 2a).

The application of overlap-type measures is popular among ecologists (e.g. for convex hulls: Braghin et al. 2013, Lindholm et al. 2020, Abedi et al. 2022; for dynamic range boxes: Cai et al. 2021, Pereira et al. 2022; for Gaussian kernels: Benavides et al. 2019, Lamanna et al. 2021, Ajal et al. 2022; for trait probability density functions: Wong et al. 2020, de Bello et al. 2021b, Carmona et al. 2021). For software availability, see the hypervolume (Blonder et al. 2022), betapart (Baselga et al. 2022), TPD (Carmona 2019) and BAT (Cardoso et al. 2022) packages.

### Discussion and Conclusions

Functional dissimilarity indices available in the literature are based on different concepts regarding the representation of communities in the trait space and dissimilarity. Modes of representation include the use of typical values, the combination of abundance and trait matrices, and hypervolumes (including several subtypes). These approaches assume increasingly complex data: for typical values, a single value per community is sufficient; species-level abundances and traits are required for the matrix approach, while individual-level trait values are ideal for hypervolumes. It is possible to apply a less data-demanding representation than the level of details available, though. For example, individual-level trait data can be arranged into abundance and trait matrices by counting the number of individuals and averaging their trait values, or community-weighted mean trait values can be derived from any type of data discussed here. Obviously, these data operations neglect some aspects of trait variation between individuals, and one must carefully measure its possible consequences on the results. Wong & Carmona (2021) argued that the level of detail, from individual-level trait measurements in each community to only species-level means across all communities, can lead to markedly different conclusions on ecological processes. However, less complex data can be tuned up to match more sophisticated requirements only at the cost of involving additional assumptions. E.g. if species-level abundances and mean trait values are available, we may assume that interspecific trait distribution is constant for all species (Lamanna et al. 2014). With this condition, it is possible to fit a TPD for all species using Gaussian kernel density estimation with fixed bandwidth and then to summarise species-level TPDs at the community level (Carmona et al. 2019).

Some of the indices were originally designed for presence-absence data (e.g. *PADDis*). Suppose species abundances are available as counts. In that case, this constraint could be resolved by taking each individual as separate “species” since this does not change the functional composition and diversity of the community, only producing a different arrangement of individuals across species. However, this does not apply to overlap indices between unweighted hypervolumes since they offer no option for weight using the density of points (but see dynamic range boxes).

Regarding the concept of dissimilarity, generally, two choices are available: distances and overlaps; however, these options are not completely distinct because some methods could be interpreted as part of both. Distances are generally less data-demanding and are suitable for detecting robust differences in the locations of the two communities in the trait space. We expect them to perform well along long gradients and strong niche filtering (Schellenberger Costa et al. 2017), even in rather complex systems (Lebrija-Trejos et al. 2010). Overlaps are more appropriate for comparing communities that are not distinct in the trait space (Mammola 2019). They seem the most straightforward when the gradient is short, and between-community variation in the shape of community-level trait distributions is also of interest. As distances and overlaps are suitable for different study situations, we reinforce Mammola (2019), who recommended trying both approaches in real study situations. In the future, the invention of a new method encompassing the positive features of distances and overlaps would be a significant addition to the field.

Notably, all methods discussed here are applicable also if species are replaced by operational taxonomical units, that can be higher or lower taxonomical units, age classes, etc. Similarly, there is no technical barrier to substitute traits with other type of information, e.g. phylogenetic relatedness or attributes describing ecological preference.

In this paper, we outlined a literature review of functional dissimilarity indices. We revealed the most fundamental decisions to consider for choosing the most suitable method.

Nevertheless, the differentiation between some alternative methods regarding sensitivity to certain biological patterns is not fully understood yet. Matrix-based approaches involve a series of abstract data operations (e.g. transformation of occurrence and trait matrices into species ordinariness or effective numbers of species, then quantification of dissimilarity), that are especially difficult to translate into biologically meaningful terms. No surprise that even the authors of the matrix-based indices pay less attention to contrast the existing and newly introduced methods from the biological viewpoint. We identify this as a major task for methodological improvements in the field of trait-based analyses. Moreover, a fully functional guidance for the users should also include the empirical performance assessment. We hope to achieve this information as well in the near future.

## Abbreviations

CWM: community-weighted mean
KDE: kernel density estimate
SVM: support vector machine

