## Appendix for "A guide to between-community functional dissimilarity measures"

**Appendix A1: Formulas of "traditional" dissimilarity indices**

In the case of presence/absence data, these indices are based on the well-known 2×2 contingency table whose cells represent the number of species shared (denoted by *a*), as well as the number of species occurring only in one of the communities (*b* and *c*). The fourth cell of the contingency table quantifying the number of shared absences is disregarded by these indices and rarely used in ecological analyses (but see Tamás et al. 2001). All these indices agree that they express similarity as the proportion of shared diversity to its possible maximum. Hence, all of them range between 0 and 1. In the case of presence/absence data the number of shared species, *a*, in the numerator stands for shared diversity for all indices, while the denominators are different. In the Sørensen index (*s_S_*) the denominator is the arithmetic mean of the species numbers of the two communities (Sørensen 1948), in Ochiai index (*s_O_*) it is their geometric mean (Ochiai 1957), in Kulczynski (*s_K_*) it is their harmonic mean (Kulczynski 1927), while in Simpson index (*s_Si_*) it is the richness of the species poorer community (Simpson 1943). If the two communities are equally species-rich, then these indices are equal, otherwise *s_S_* < *s_O_* < *s_K_* < *s_Si_*. In the Jaccard index (*s_J_*), the denominator is the total number of species in the two communities (Jaccard 1901), while in Sokal & Sneath index (*s_SS_*) species occurring in a single community are taken into account with double weight (Sokal & Sneath 1963). There is a direct and monotonic relationship between Jaccard, Sørensen, and Sokal & Sneath indices (see Appendix A2). Table A1 summarises the similarity and dissimilarity forms of the above indices.

For abundance data, the properties and logic of indices are discussed in the main text. Here only the formulas are given. Let us denote the abundances of species *i* in communities *j* and *k*, by *x_ij_* and *x_ik_,* and the total number of species in *j* and *k* by *S_jk_*.

Euclidean distance:

Eq. A1-1.	$d_{Euclidean}=\sqrt{\sum_{i=1}^{S_{jk}} \left( x_{ij}-x_{ik} \right)^{2}}$

Manhattan distance:

Eq. A1-2.	$d_{Manhattan}=\sum_{i=1}^{S_{jk}} \left| x_{ij}-x_{ik} \right|$

Canberra index (Lance & Williams 1966):

Eq. A1-3.	$d_{Canberra}=\sum_{i=1}^{S_{jk}} \frac{\left| x_{ij}-x_{ik} \right|}{\left( x_{ij}+x_{ik} \right)}$

normalised Canberra index (Ricotta & Podani 2017):

Eq. A1-4.	$d_{NCanberra}=\frac{1}{S_{jk}}\sum_{i=1}^{S_{jk}} \frac{\left| x_{ij}-x_{ik} \right|}{\left( x_{ij}+x_{ik} \right)}$

normalised modified Canberra index (Ricotta & Podani 2017):

Eq. A1-5.	$d_{NMCanberra}=\frac{1}{S_{jk}}\sum_{i=1}^{S_{jk}} \frac{\left| x_{ij}-x_{ik} \right|}{\max_{} (x_{ij},x_{ik})}$

Bray-Curtis index (Bray & Curtis 1957):

Eq. A1-6.	$d_{BC}=\frac{\sum_{i=1}^{S_{jk}} \left| x_{ij}-x_{ik} \right|}{\sum_{i=1}^{S_{jk}} (x_{ij}+x_{ik})}$

Marczewski-Steinhaus index (Marczewski & Steinhaus 1958):

Eq. A1-7.	$d_{MS}=\frac{\sum_{i=1}^{S_{jk}} \left| x_{ij}-x_{ik} \right|}{\sum_{i=1}^{S_{jk}} \max_{} (x_{ij},x_{ik})}$

Several abundance-based indices can be expressed if we generalise the *a*, *b*, and *c* quantities used during the definition of indices for presence/absence data (Tamás et al. 2001).

Eq. A1-8.	$a^{'}=\sum_{i=1}^{S_{jk}} \min_{} \left( x_{ij},x_{ik} \right)$

Eq. A1-9.	$b^{'}=\sum_{i=1}^{S_{jk}} \left( \max_{} \left( x_{ij},x_{ik} \right)-x_{ij} \right)$

Eq. A1-10.	$c^{'}=\sum_{i=1}^{S_{jk}} \left( \max_{} \left( x_{ij},x_{ik} \right)-x_{ik} \right)$

Substituting *a*, *b* and *c* with *a'*, *b'* and *c'* into the formula of the Sørensen index gives Bray-Curtis and doing so with Jaccard index results in the Marczewski-Steinhaus. Abundance versions of all other presence/absence indices can be created in the same manner.

The general formula for (niche) overlap proposed by Horn (1966) is the following:

Eq. A1-11.	$O_{jk}=\frac{H_{max}-H_{jk}}{H_{max}-H_{min}}$

where H_jk_ is the diversity of the pooled sample, while H_max_ and H_min_ are its possible maximum and minimum, respectively. For Shannon diversity

Eq. A1-12.	$H_{jk}=-\sum\frac{x_{ij}+x_{ik}}{x_{.j}+x_{.k}}\log\frac{x_{ij}+x_{ik}}{x_{.j}+x_{.k}}=\log\left( x_{.j}+x_{.k} \right)-\frac{1}{x_{.j}+x_{.k}}\sum\left( x_{ij}+x_{ik} \right)\log\left( x_{ij}+x_{ik} \right)$

The pooled diversity is maximal if no shared species:

Eq. A1-13.	$H_{max}=\log\left( x_{.j}+x_{.k} \right)-\frac{1}{x_{.j}+x_{.k}}\left[ \sum x_{ij}\log x_{ij}+\sum x_{ik}\log x_{ik} \right]$

while the possible minimum is the weighted mean of Shannon entropy in the two samples:

Eq. A1-14.	$H_{min}=\frac{x_{.j}}{x_{.j}+x_{.k}}\left( \log x_{.j}-\frac{1}{x_{.j}}\sum x_{ij}\log x_{ij} \right)+\frac{x_{.k}}{x_{.j}+x_{.k}}\left( \log x_{.k}-\frac{1}{x_{.k}}\sum x_{ik}\log x_{ik} \right)=\frac{1}{x_{.j}+x_{.k}}\left[ x_{.j}\log x_{.j}+x_{.k}\log x_{.k}-\sum x_{ij}\log x_{ij}-\sum x_{ik}\log x_{ik} \right]$

Substituting them to the general formula,

Eq. A1-15.	$O_{jk}=\frac{\sum\left( x_{ij}+x_{ik} \right)\log\left( x_{ij}+x_{ik} \right)-\sum x_{ij}\log x_{ij}-\sum x_{ik}\log x_{ik}}{\left( x_{.j}+x_{.k} \right)\log\left( x_{.j}+x_{.k} \right)-x_{.j}\log x_{.j}-x_{.k}\log x_{.k}}$

Let us calculate the overlap between two communities which differ only in total abundance, but have the same relative abundances, i.e. *x_ik_=cx_ij_.* In this case the numerator is

Eq. A1-16.	$\sum\left( c+1 \right)x_{ij}\log\left[ \left( c+1 \right)x_{ij} \right]-\sum x_{ij}\log x_{ij}-\sum cx_{ij}\log{cx}_{ij}=\left( c+1 \right)\sum x_{ij}\log x_{ij}+\left( c+1 \right)\sum x_{ij}\log\left( c+1 \right)-\sum x_{ij}\log x_{ij}-c\sum x_{ij}\log x_{ij}-c\sum x_{ij}\log c=\left( c+1 \right)x_{.j}\log\left( c+1 \right)-cx_{.j}\log c$

while the denominator is

Eq. A1-17.	$\left( c+1 \right)x_{.j}\log\left( c+1 \right)x_{.j}-x_{.j}\log x_{.j}-{cx}_{.j}\mathrm{logc} x_{.j}=\left( c+1 \right)x_{.j}\log\left( c+1 \right)-cx_{.j}\log c$

thus the overlap (i.e. similarity) is 1. It means that Horn index is sensitive only to relative abundances, and insensitive to total community size.

For deducing overlap from Gini-Simpson diversity, let us suppose that *x_ij_* is a relative abundance, thus $\sum_{i} x_{ij}=1$. In this case, the relative abundances in the pooled sample are the unweighted means of the relative abundances in the two compared samples and diversity of the pooled samples is

Eq. A1-18.	$H_{ij}=1-\sum\frac{\left( x_{ij}+x_{ik} \right)^{2}}{4}=1-\frac{1}{4}\sum\left( x_{ij}^{2}+x_{ik}^{2}+2x_{ij}x_{ik} \right)$

Again, it is maximal if no shared species:

Eq. A1-19.	$H_{max}=1-\frac{1}{4}\sum\left( x_{ij}^{2}+x_{ik}^{2} \right)$

and its possible minimum is the unweighted mean of the diversity in the two samples:

Eq. A1-20.	$H_{min}=1-\frac{1}{2}\sum\left( x_{ij}^{2}+x_{ik}^{2} \right)$

Substituting to the general formula lead to the Morisita-Horn index:

Eq. A1-21.	$O_{jk}=\frac{2\sum x_{ij}x_{ik}}{\sum\left( x_{ij}^{2}+x_{ik}^{2} \right)}$

Note that if xij is absolute, instead of relative abundance, the Morisita-Horn index becomes sensitive to differences in total abundances. If two compared communities differ only in total abundances (i.e. *x_ik_=cx_ij_*) their Morisita-Horn similarity is

Eq. A1-22.	$O_{jk}=\frac{2c}{c^{2}+1}$

thus Morisita-Horn similarity calculated from raw (absolute) abundances is sensitive to differences in community size.

Diversity can be calculated for vector of abundances of a given species across communities. Dividing diversity by its possible maximum is one, if species' abundance is the same in all communities, and zero if it occurs only in one community. Ricotta (2018) proposed that this evenness can be regarded as species' contribution to the similarity, and their weighted sum can be used as similarity a measure.

Note, that the last three indices (Horn, Morisita-Horn and Ricotta's evenness based similarity) easily can be generalised for multi-site situations.

Table A1. Similarity and dissimilarity forms of resemblance indices for presence-absence data. Notations: *k*, *j* – two communities under comparison; *Sj*, *Sk* – total species numbers of communities *j* and *k*; *a* – number of shared species; *b*, *c* – number of species present either in community *j* or in community *k*; *d* – number of species absent in *j* and *k*.

| Name of the index | Similarity version | Dissimilarity version |
| --- | --- | --- |
| Sørensen | $s_{S}=\frac{2a}{2a+b+c}=\frac{a}{\left( S_{j}+S_{k} \right)/2}$ | $d_{S}=\frac{b+c}{2a+b+c}=\frac{b+c}{S_{j}+S_{k}}$ |
| Ochiai | $s_{O}=\frac{a}{\sqrt{\left( a+b \right)\left( a+c \right)}}=\frac{a}{\sqrt{S_{j}S_{k}}}$ | $d_{O}=\frac{b+c}{\sqrt{S_{j}S_{k}}}$ |
| Kulczynski | $s_{K}=\frac{1}{2}\left( \frac{a}{a+b}+\frac{a}{a+c} \right)=\frac{a}{2/\left( 1/{S_{j}}+1/{S_{k}} \right)}$ | $d_{K}=\frac{1}{2}\left( \frac{b}{a+b}+\frac{c}{a+c} \right)=\frac{1}{2}\left( \frac{b}{S_{j}}+\frac{c}{S_{k}} \right)$ |
| Simpson | $s_{Si}=\frac{a}{a+\min_{} (b, c)}=\frac{a}{\min_{} \left( S_{j}, S_{k} \right)}$ | $d_{Si}=\frac{b+c}{\min_{} (S_{j},S_{k})}$ |
| Jaccard | $s_{J}=\frac{a}{a+b+c}=\frac{a}{S_{jk}}$ | $d_{J}=\frac{b+c}{a+b+c}=\frac{b+c}{S_{jk}}$ |
| Sokal & Sneath | $s_{SS}=\frac{a}{a+2\left( b+c \right)}$ | $d_{SS}=\frac{2\left( b+c \right)}{a+2\left( b+c \right)}$ |

**Appendix A2. Algebraic relationship of Jaccard, Sörensen and Sokal-Sneath indices**

Jaccard (*S_J_*), Sörensen (*S_S_*) and Sokal-Sneath (*S_SS_*) indices are special cases of the following general formula with parameters *v>0* and *z>0*:

Eq. A2-1.	$S_{vz}=\frac{va}{va+z(b+c)}$

*a*: number of species shared between community 1 and community 2

*b*: number of species present only in community 1

*c*: number of species present only in community 2

Any member of this family can be expressed as the monotonic transformation of Jaccard index:

Eq. A2-2.	$S_{vz}=\frac{vS_{J}}{\left( v-z \right)S_{J}+z}$

Proof

Eq. A2-3	.$s_{J}=\frac{a}{a+b+c}$

Eq. A2-4.	$\frac{1}{S_{J}}=1+\frac{b+c}{a}$

Eq. A2-5	.$\frac{b+c}{a}=\frac{1}{S_{j}}-1=\frac{1-S_{j}}{S_{j}}$

Eq. A2-6.	$\frac{1}{S_{vz}}=1+\frac{z}{v}*\frac{b+c}{a}=1+\frac{z}{v}*\frac{1-S_{J}}{S_{J}}=\frac{vS_{J}+z\left( 1-S_{J} \right)}{vS_{J}}=\frac{\left( v-z \right)S_{J}+z}{vS_{J}}$

Eq. A2-7	.$S_{vz}=\frac{vS_{J}}{\left( v-z \right)S_{J}+z}$

Sörensen index can be obtained with v=2 and z=1:

Eq. A2-8.	$S_{S}=\frac{2S_{J}}{S_{J}+1}=\frac{\frac{2a}{a+b+c}}{\frac{a+a+b+c}{a+b+c}}=\frac{2a}{2a+b+c}$

Sokal-Sneath index can be derived with v=1 and z=2:

Eq. A2-9	.$S_{SS}=\frac{S_{J}}{2-S_{J}}=\frac{\frac{a}{a+b+c}}{\frac{2\left( a+b+c \right)-a}{a+b+c}}=\frac{a}{a-2b-2c}$

See also Janson & Vegelius (1981), Hubálek (1982).
